# E-cadherin is a structuring component of invadopodia in pancreatic cancer

**DOI:** 10.1101/2020.10.09.332783

**Authors:** Aurélie Dobric, Sébastien Germain, Françoise Silvy, Rénaté Bonier, Stéphane Audebert, Luc Camoin, Nelson Dusetti, Philippe Soubeyran, Juan Iovanna, Véronique Rigot, Frédéric André

**Author notes:** Corresponding authors: Frédéric ANDRE, Véronique RIGOT. both authors contribute equaly to the work (Françoise Silvy, Rénaté Bonier).

## Abstract

**Graphical abstract:** 

**Background:** The appearance of hybrid epithelial-mesenchymal (E/M) cells expressing E-cadherin is favourable for the establishment of pro-invasive function. However, the molecular mechanism and potential roles of E-cadherin in cancer cell invasion stay unexplored.

**Methods:** We used models of E/M hybrid cell lines, tissues sections and patient-derived xenografts from a multi-center clinical trial. E-cadherin involvement in invadopodia formation was assessed using a gelatin-FITC degradation assay. Mechanistic studies were performed by using proteomic analysis, siRNA strategy and proximity ligation assay.

**Results:** We showed that E-cadherin is a critical component of invadopodia. This unexpected localization results from a synergistic trafficking of E-cadherin and MT1-MMP through Rab vesicle-dependent pathway. Modulation of E-cadherin expression or activation impacted invadopodia formation. <u>Moreover, colocalization of E-cadherin and Actin in ―ring structures‖</u> <u>as precursor of invadopodia reveals that E-cadherin is required for invadopodia structuration.</u>

**Conclusion:** E-cadherin, initially localized in the adherens junctions could be recycled to nascent invadopodia where it will interact with several components such as Arp2/3, Cortactin or MT1-MMP. <u>The trans-adhesive properties of E-cadherin are therefore essential for structuring</u> <u>invadopodia</u>.

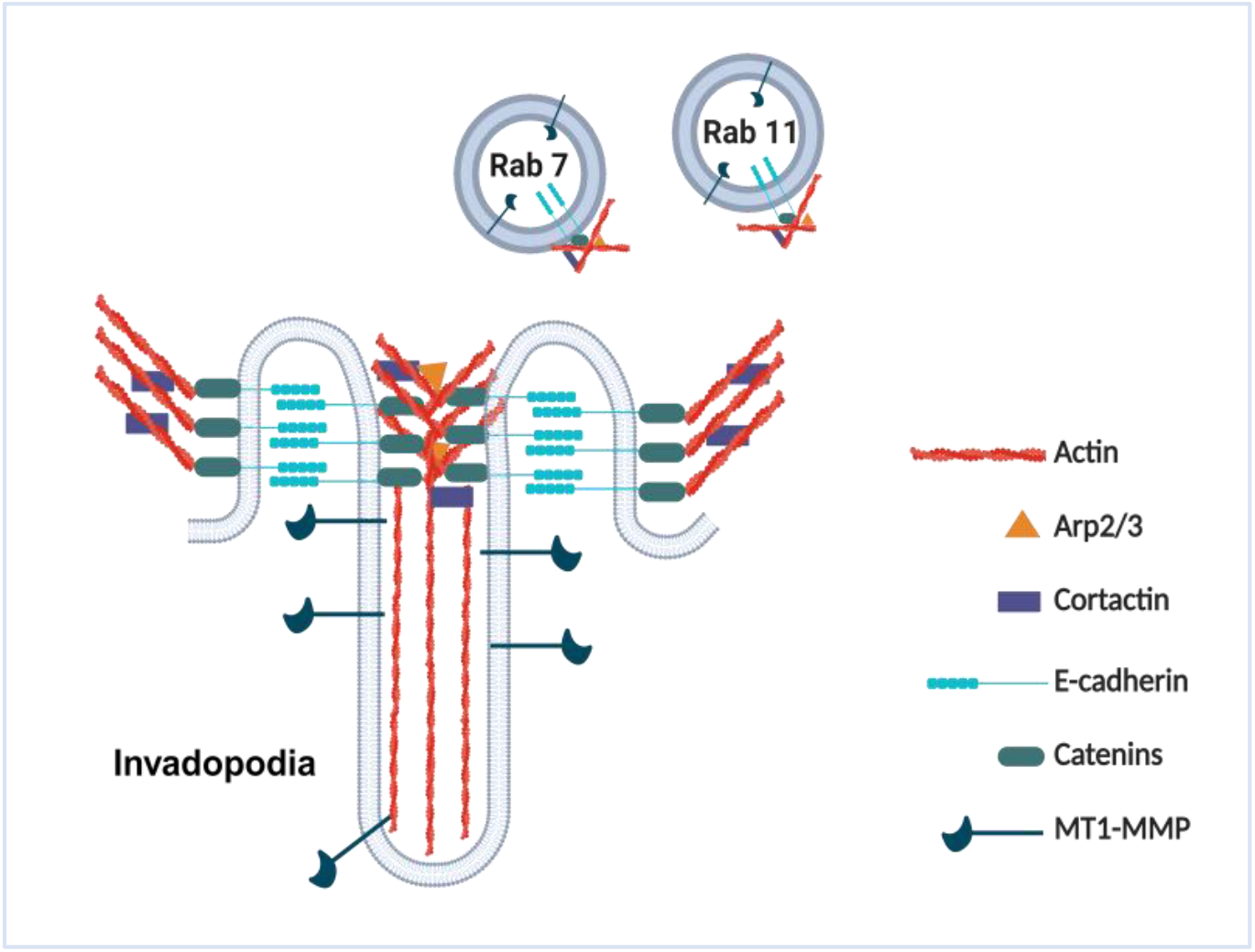

## Background

The epithelial-to-mesenchymal transition (EMT) is a key biological process associated with the gain, either individually or collectively, of mesenchymal features and the acquisition of migration and invasion properties by cancer cells, conferring metastasis properties (Dongre & Weinberg, 2019). During this process, epithelial cells lose their apical-basal polarity, remodel their cytoskeleton and exhibit reduced cell-cell adhesion properties (Yang *et al*, 2020).

Studies have shown that EMT is a process with distinct intermediates, reflecting a progressive acquisition and a loss of mesenchymal and epithelial molecular traits. Such traits coexists in intermediates states, as documented by a mixture of epithelial and mesenchymal features at molecular and morphological levels (Pastushenko *et al*, 2018; Jolly *et al*, 2019; Kröger *et al*, 2019). These « multiple shadows » mirror hybrid epithelial/mesenchymal (E/M) phenotypes of cells, with distinct biological properties (Pastushenko *et al*, 2018). In cancer, a hybrid E/M signature contributes to an intra-tumoural heterogeneity and is an indicator of poor prognosis, as cancer cells may metastasize with a partial loss of epithelial and a partial gain of mesenchymal traits (Andriani *et al*, 2016; Saitoh, 2018; Jolly *et al*, 2019; Simeonov *et al*, 2021; Canciello *et al*, 2022).

Classical cadherins establish adhesion between neighbouring cells through their extracellular domain and ensure the cohesion required for tissue integrity (Niessen *et al*, 2011). Their intracellular domain is associated with catenins, which allow connection to the actin cytoskeleton and cell signalling pathways. Complete downregulation of epithelial E-cadherin associated with an up-regulation of N-cadherin and/or P-cadherin during the EMT process was historically considered a crucial step in carcinoma progression to promote invasion and metastasis (Thiery *et al*, 2009). However, a correlation between low E-cadherin expression, cell invasion and metastasis is not absolute. Indeed, studies have shown than tumour cells with epithelial traits and still expressing E-cadherin can undergo metastasis and form secondary tumours (Reddy *et al*, 2005; Lewis-Tuffin *et al*, 2010; Putzke *et al*, 2011; Sulaiman *et al*, 2018; Sommariva & Gagliano, 2020; Canciello *et al*, 2022). Moreover, E-cadherin was reported as a promoter of metastasis in models of invasive ductal breast carcinomas (Padmanaban *et al*, 2019; Shen & Kang, 2019). However, the molecular mechanisms involved in this process remain to be elucidated.

During metastasis progression, invading cells acquire the capability to degrade the extracellular matrix and ultimately invade the vasculature. These processes are driven by invadopodia-actin rich protrusive plasma membrane structures that operate focalized proteolysis (Paterson & Courtneidge, 2018; Ferrari *et al*, 2019; Linder *et al*, 2023).

Key components of invadopodia include the scaffold protein Tks5, the actin regulators Cortactin, Wiskott–Aldrich syndrome protein family members, cofilin, and membrane type 1 matrix metalloproteinase (MT1-MMP) (Paterson & Courtneidge, 2018). Invadopodia have been extensively studied in cell culture, and have been detected in vivo (Génot & Gligorijevic, 2014; Chen *et al*, 2019). Whereas significant advances have been made in understanding how invadopodia formation and activity are regulated, a putative role of cadherins in invadopodia organization remains unknown.

Pancreatic ductal adenocarcinoma (PDAC) is a cancer with poor prognosis (Siegel *et al*, 2022). This aggressivity is due to a combination of factors including a lack of early diagnostic markers, lack of symptoms and an early metastatic spread. The desmoplastic reaction observed in PDAC is a hallmark of disease progression and prognosis. It is correlated with inflammation and a low vascularity. The presence of large amounts of extracellular matrix (ECM) components and tumour-infiltrating leukocytes are determinant in EMT evolution and therapeutic resistance (Beatty *et al*, 2021; Bhoopathi *et al*, 2023).

PDAC encompasses a range of E/M hybrid cells, reflecting epithelial-mesenchymal plasticity (Andriani *et al*, 2016; Saitoh, 2018; Jolly *et al*, 2019; Simeonov *et al*, 2021). Intriguingly, it has been shown that some PDAC cells with high E-cadherin expression at the cell boundaries exhibit highly invasive and malignant behaviour (Sommariva & Gagliano, 2020). PDAC represents therefore an appropriate biological context to explore cadherins implication in invasive front migration and capabilities to form invadopodia.

The aim of this study is to decipher how in pancreatic cancer, E-cadherin modulates the ability of E/M hybrid cells to degrade ECM.

## Methods

### Antibodies and reagents

Mouse anti-E-cadherin (M168 and HECD-1), rabbit anti-E-cadherin (EP700Y) and anti-Arp3 (EPR110429) were from Abcam. Mouse anti-E-cadherin (24E10), rabbit anti-P-cadherin (21300S) and rabbit anti-Rab7 (D95F2) were from Cell Signaling. Mouse anti-Cortactin p80/p85 (4F11) and anti-MT1-MMP (LEM 2/15.8) were from Millipore. Goat anti-E-cadherin was from St John‘s Laboratory. Rabbit anti-Rab11 and rhodamine-conjugated phalloidin were from Life Technologies. Rabbit anti Tks5 was from Novus. Alexafluor 488, 594, 647 secondary antibodies were from Thermo Fisher. Both AS9 (BAS00132635) and AS11 (BAS00602705) compounds were from Asinex.

### Cell culture

The human pancreatic adenocarcinoma BxPC-3 cells, authenticated using short tandem repeat (STR) profiling (ATCC), were cultured as previously published (Siret *et al*, 2018). E-cadherin was stably knocked down using shRNA lentiviral transduction particles as previously described (Siret *et al*, 2018). Primary cell cultures PDAC001T and PDAC021T derived from patient-derived xenograft (PDX) and SUM 149 cell line derived from inflammatory breast cancer were cultured as previously described, respectively (Fabre *et al*, 1993; Hoffmeyer *et al*, 2005; Siret *et al*, 2018) <u>E-cadherin deficient PDAC021T were</u> <u>transfected with human E-Cadherin mGFP-tagged Tagged ORF Clone Lentiviral Particle</u> <u>(Origene) at 25 multiplicity of infection (MOI). Infected cells were selected using 2,5μg/ml</u> <u>puromycin. E-cadherin expression was checked by western blot analysis.</u>

### Reverse siRNA transfection

ON-target plus smartpool human control, human MT1-MMP, human Arp3, human Rab7A and human Rab11A were from Dharmacon. Cells were seeded in 6-well plates directly with the siRNA/transfection mix: 3µl of LipoRNAiMax (Life Technologies), 500µL of OptiMEM (Life Technologies), 25 or 50nM of indicated siRNA and 2.5ml of RPMI/10% FCS medium. When required, transfected cells were detached and seeded on FITC-gelatin coated coverslips, 24 or 48 h after treatment.

### Subcutaneous xenografts of pancreatic cancer cells

All experimental procedures involving animals were performed in accordance with French Guidelines and approved by the ethical committee of Marseille (agreement 50-31102012). BxPC-3 cells were harvested by mild trypsinisation, washed twice in PBS, then suspended in Matrigel at 2×10^6^ cells per 100 μl. To induce tumours, the cell suspension was injected subcutaneously (s.c.) into the flank of 6–8-week-old female NMRI-Foxn1nu/Foxn1nu mice (Charles River Laboratories, L‘Arbresle, France). Mice were sacrificed 3 weeks after inoculation. Tumours were removed and tissue specimens were fixed in 4% formalin then embedded in paraffin.

### Immunohistofluorescence

Human pancreatic cancer samples were obtained as previously described (approval DC2013-1857) (Martinez *et al*, 2016). Human tissue specimens or mouse tumours were cut into 3 µm sections. After dewaxing and antigen retrieval at pH9, sections were incubated for double or triple staining with primary antibodies for 2 h at room temperature. After washing, the sections were incubated with Alexa Fluor-conjugated antibodies, washed, and mounted in aqueous mounting medium. Images were captured with an LSM 880 Zeiss confocal microscope equipped with ZEN Software (objective 40X). <u>Colocalization quantifications were performed using Jacop plugging (FiJi software). Overlap</u> <u>coefficients were obtained by dividing the number of points in the overlap region (different channels)</u> <u>with the total numbers of points in one of the distributions (each channel).</u>

### Indirect immunofluorescence microscopy

Cells were fixed in 4% formaldehyde for 30 min then permeabilized and blocked with phosphate buffered saline/bovine serum albumin (PBS/BSA) 4% saponin 0.1% for 1 h. Cells were successively incubated with indicated primary antibodies in PBS/BSA 1% saponin 0.1% for 2 h at RT and with Alexa Fluor-conjugated secondary antibodies in PBS/BSA 1% saponin 0.1% for 1 h raised against mouse or rabbit immunoglobulins (Invitrogen). After washes, samples were mounted in ProLong Gold antifade reagent (Thermo Fisher Scientific). Images were acquired with an Sp5 Leica or LSM 880 Zeiss confocal microscopes equipped, respectively, with LAS AF Lite or ZEN Software. Z-Stack acquisitions (Range: 0.5µm) were performed using a 63X objective magnification and analysed through orthogonal projections using ImageJ software (<u>rsb.info.nih.gov/ij/</u>). <u>Actin and E-cadherin ring structures were obtained using the ZEIS</u> <u>LSM880 AiryScan 2.5D module.</u>

### Invadopodia assay

Coverslips were coated with FITC-conjugated gelatin (Life Technologies), fixed with 0.5% glutaraldehyde and incubated for 3 min at RT with 5 mg/ml sodium borohydride (Sigma). After washes, 10^4^ isolated cells were seeded on top of the coverslip. Cells were incubated 16 h at 37°C, fixed in 4% formaldehyde and stained for proteins of interest as described previously. The areas of degraded matrix were observed using a LSM880 Zeiss confocal microscope (20X objective). 15 microscopic fields per coverslip were acquired with all fluorescent channels.

### Kinetic of invadopodia formation

Lab-Teck chamber coverglasses were coated as described for invadopodia assays. 2 hours after seeding, cells were incubated for 18h with AS11 (0.01 mM) or DMSO in a temperature and CO_2_ controlled chamber mounted on an Olympus IX83 inverted microscope. Cells were then washed and incubated in DMEM/10% FBS for an additional 24h period. Invadopodia formation was analysed by videomicroscopy by capturing images every hour using an orca-flash4 camera with a 40X objective.

### Immunoprecipitation

BxPC-3 cells were plated in 10 cm^2^ culture dishes. Subconfluent cells (70% of confluence) were lyzed in ice with lysis buffer (50mM HEPES pH 7.5; 150 mM NaCl; 1 mM EDTA; 1 mM EGTA; Glycerol 10%; Triton X-100 1%; 25 mM NaF; 10 µM ZnCl_2_ + protease inhibitor cocktail). Protein G Sepharose beads (Roche) were pre-incubated with 1 µg of indicated primary antibody for 2 h at 4°C. After washes, equal amounts of cell lysate were incubated with pre-incubated beads for 2 h at 4°C. After three washes in PBS, immunoprecipitated proteins were solubilized in Laemmli buffer, heated at 100°C for 5 min, and analysed by western blotting.

### Western Blotting

Cells were lyzed with 150mM RIPA Buffer (25mM Tris-HCl pH 8.0; 150mM NaCl; 1% Triton-X100) containing protease inhibitor cocktail. Equal amounts of cell lysate (25µg) were resolved by SDS PAGE (8 or 10% polyacrylamide) and blotted onto a polyvinylidene difluoride (PVDF) membrane. Proteins were detected using indicated antibodies. Antigen–antibody complexes were revealed using the ECL detection system (Millipore) and detected using a Pxi imaging device (SynGene).

### Invadopodia fractioning

The isolation of an enriched fraction of invadopodia was performed using previously published protocol (Attanasio *et al*, 2011). Cells were seeded at 2.5×10^5^ cells on 4 culture dishes (10 cm diameter) coated with non-fluorescent gelatin. After 18 h, plates were washed in PBS containing 0.5 mM MgCl2, 1 mM CaCl2, then in five times diluted PBS containing 0.5 mM MgCl2, 1 mM CaCl2, and incubated for 15 min in the presence of 3 ml of the diluted PBS containing protease inhibitor mixture to induce cell swelling. Cell bodies were then sheared away using an L shaped Pasteur pipette with sealed end, to leave invadopodia embedded in the gelatin. The embedded invadopodia were then washed in PBS containing 0.5 mM MgCl2, 1 mM CaCl2 until no cell body were visible on the dishes. Then the embedded invadopodia were scraped away with the gelatin into lysis buffer (150 mM NaCl, 1% NP40, 0,5% sodium deoxycholate, 0,1% sodium dodecyl sulphate, 50 mM Tris base buffer pH 8, proteases inhibitor) and clarified by centrifugation (15 min, 13,000 rpm at 4°C). The cell body fraction was further separated into cell body membranes and cytosol fractions by centrifugation at 9,000 g for 20 min at 4°C. The supernatant (cytosolic fraction) was discarded whereas the cell body membrane pellet obtained after centrifugation was solubilized in lysis buffer and clarified by centrifugation (15 min, 13,000 rpm at 4°C). Both invadopodia and cell membrane fractions were precipitated in 3 volumes of cold acetone overnight at −20°C, centrifuged and denatured. All the invadopodia membrane fraction corresponding for 4 dishes was loaded with 1/2 of cell body membrane fraction.

### Proximity ligation assay

Proximity ligation assay (PLA) was performed according to the manufacturer‘s recommendations protocol (Duolink; Sigma). Briefly, cells were prepared as for indirect immunofluorescence. Cells were incubated with indicated primary antibodies for 2h at RT. After washing, samples were incubated with the respective PLA probes (Duolink in situ probes anti-Rb PLUS and Duolink in situ probe anti-Mouse MINUS) for 1 h at 37°C, washed and then ligated for 30 min at 37°C. Amplification with polymerase was then performed for 100 min at 37°C in the dark. After washes, nuclei were stained with DAPI, and samples are mounted in ProLong Gold antifade reagent. Images were captured as described in the indirect immunofluorescence staining section.

### Mass spectrometry analysis

Proteomic analysis from E-cadherin depleted cells (BxPC-3 shEcad) were compared to control cells (BxPC-3 shCTRL) by label-free quantitative mass spectrometry analysis. Briefly we used 15 µg of each cell lysate for proceeding and trypsin digestion (Shevchenko *et al*, 1996). Details of samples preparation and data processing protocols (Perez-Riverol *et al*, 2019) are available in **additional data 1**.

### Pathway enrichment analysis

Proteins identified by mass spectrometry were analysed with Ingenuity Pathway Analysis (IPA) software to study pathway enrichment. The statistical significance of the enrichment was calculated using FDR method (*P*-value < 0.05). Protein Z-score was calculated for each protein of the selected pathway. Z-score indicate the overall activation state.

### Statistics

Data are presented as the mean ± SEM for three independent experiments performed in triplicate. Comparison between two conditions was made using the Mann– Witney test: *P* < 0.05 was considered statistically significant in all analyses and is indicated by ―***‖ when *P* < 0.001, ―**‖ when *P* < 0.01 and ―*‖ when *P* < 0.05.

## Results

### E-cadherin localizes within invadopodia

We previously described that both E-cadherin and P-cadherin are jointly expressed at the cell surface of tumoural cells in a large proportion of PDAC, pointing to the importance of hybrid E/M cells expressing E-cadherin in this pathology (Siret *et al*, 2018). Previous studies have shown that E-cadherin at the cell boundaries exhibits highly invasive and malignant behaviour (Sommariva & Gagliano, 2020). To strengthen this function in pancreatic cancer, we used *in vitro* approaches to determine how E-cadherin regulates cell invasion by analysing the formation of invadopodia, an early step of the invasion process.

The hybrid E/M BxPC-3 cell model was first used since it express high levels of E-cadherin (Siret *et al*, 2018). X-Z confocal projections showed localization of actin spots with several invadopodia markers (Cortactin, Tks5 and MT1-MMP) within a degradation area of the FITC-labelled gelatin ***(Fig.S1A, S1B & S1C)***. Moreover, MMPs inhibitors ***(Fig.S1D*** *and **S1E)*** and MT1-MMP depletion by siRNA strategy *(**Fig. S1F** and **S1G**)* almost entirely reduced both the capacity of these cells to degrade gelatin and the number of cells forming invadopodia. MT-MMP1 depletion using siRNA strategy was controlled by western blot ***(Fig.S1H)***. This indicates that in hybrid E/M BxPC-3 cell model exhibits active invadopodia.

Immunostaining analysis, using antibodies raised either against the intracellular ***(Fig.1A***) or the extracellular domain of E-cadherin ***(Fig.1B)*** revealed that a pool of E-cadherin is located at the invadopodial membrane. P-cadherin, also expressed by BxPC-3 cells (Siret *et al*, 2018), was not detected at the invadopodial membrane ***(Fig.1C)***, indicating a specificity of E-cadherin localization in invadopodia. <u>Biochemical analysis on fraction enriched in</u> <u>invadopodia reassuringly confirmed the presence in the invasive structure of E-cadherin, in</u> <u>association with β-catenin. As observed in immunostaining analysis, P-cadherin is not</u> <u>detected in the invadopodia fraction by western-blot and could be considered as a negative</u> <u>control of cell membrane contamination. Histone H1 is not detected in invadopodia fraction</u> <u>as negative control for cell body contamination</u> ***(Fig.1D)***.

**Fig.1:**
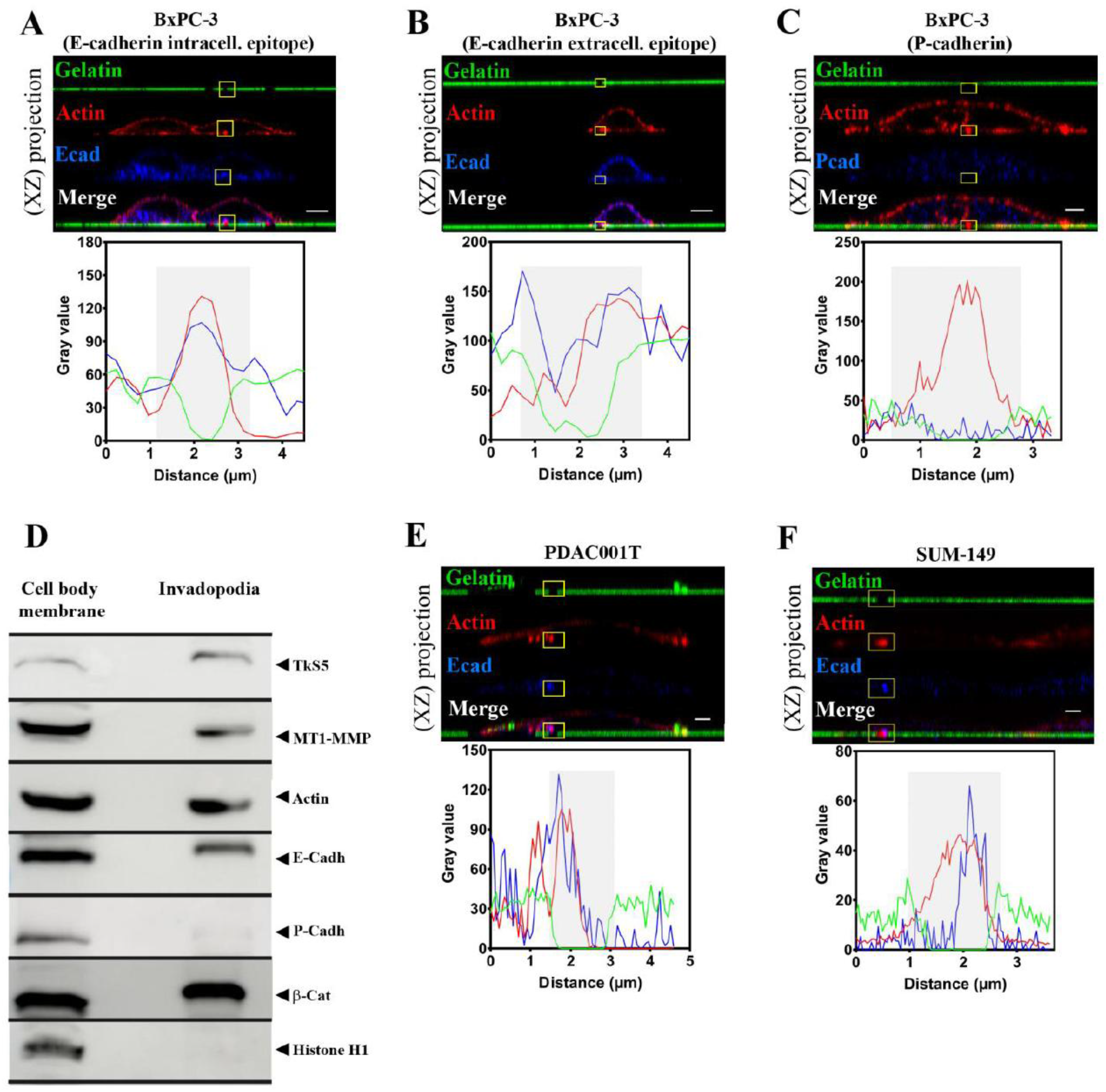
E-cadherin localizes within invadopodia. Pancreatic cancer BxPC-3 cell line **(A-C)**, pancreatic cancer primary culture PDAC001T **(E)** and breast cancer cells SUM-149 cell line **(F)** were cultured on FITC-labelled gelatin. Cells were stained for actin with phalloidin-rhodamin (red) and E-cadherin using an antibody raised against the cytoplasmic domain (blue) **(A)** or extracellular domain **(B)** or P-cadherin **(C)**. An actin spot localization with a degradation zone of the FITC-labelled gelatin represents an active invadopodia. Top panel: Images represent Z-stack confocal acquisitions. Scale bar = 10 µm **(A, B)** or 2 µm **(C, E, F)**. Bottom panel: fluorescence intensity quantification of the region of interest indicated by the yellow square on the top panel. The gelatin degradation area is identified in grey. **(D)** The BxPC-3 cell body membrane and invadopodia membrane were enriched as described in Methods, subjected to SDS-PAGE and transferred onto nitro-cellulose membrane. TkS5, E-cadherin, MT1-MMP, P-cadherin, Actin and Histone H1 were sequentially detected by western blot in the same membrane. Images in 2D view for (**A**) and (**C**) are available in ***Fig.S4A*. (A):** A representative image of 7 experiments with 5 acquisitions for each (n=7), **(B-C):** A representative image of 3 experiments with 5 acquisitions for each (n=3), **(D):** One experiment representative of 3, **(E-F):** A representative image of 2 experiments with 5 acquisitions for each (n=2).

E-cadherin was also detected in invadopodia of primary pancreatic cancer cells PDAC001T ***(Fig.1E)*** and SUM-149 cell line derived from inflammatory breast cancer ***(Fig.1F)***. Therefore, the localization of E-cadherin in invadopodia could be extended to other cell and cancer types.

### E-cadherin localizes with Cortactin and Tks5 at invadopodia-like-structures in PDAC

To confirm these observations in vivo, we immunostained E-cadherin on patient tissues, as well as Cortactin and Tks5. A Cortactin/Tks5/E-cadherin triple staining on tissue sections revealed that a pool of E-cadherin localizes with Cortactin and Tks5 at plasma membrane of cells localized in the contact with extracellular matrix ***(Fig. 2A)***. <u>For each panel we quantified</u> <u>the overlap coefficients: for E-cadherin/cortactin (75.8%), E-cadherin/TkS5 (84.6%) and</u> <u>Tks5/cortactin (70.3%).</u> This triple Cortactin/Tks5/E-cadherin colocalization was observed in 3 out of 6 patient tissues indicating that invadopodia-like structures could be detected in patient tissues.

**Fig.2:**
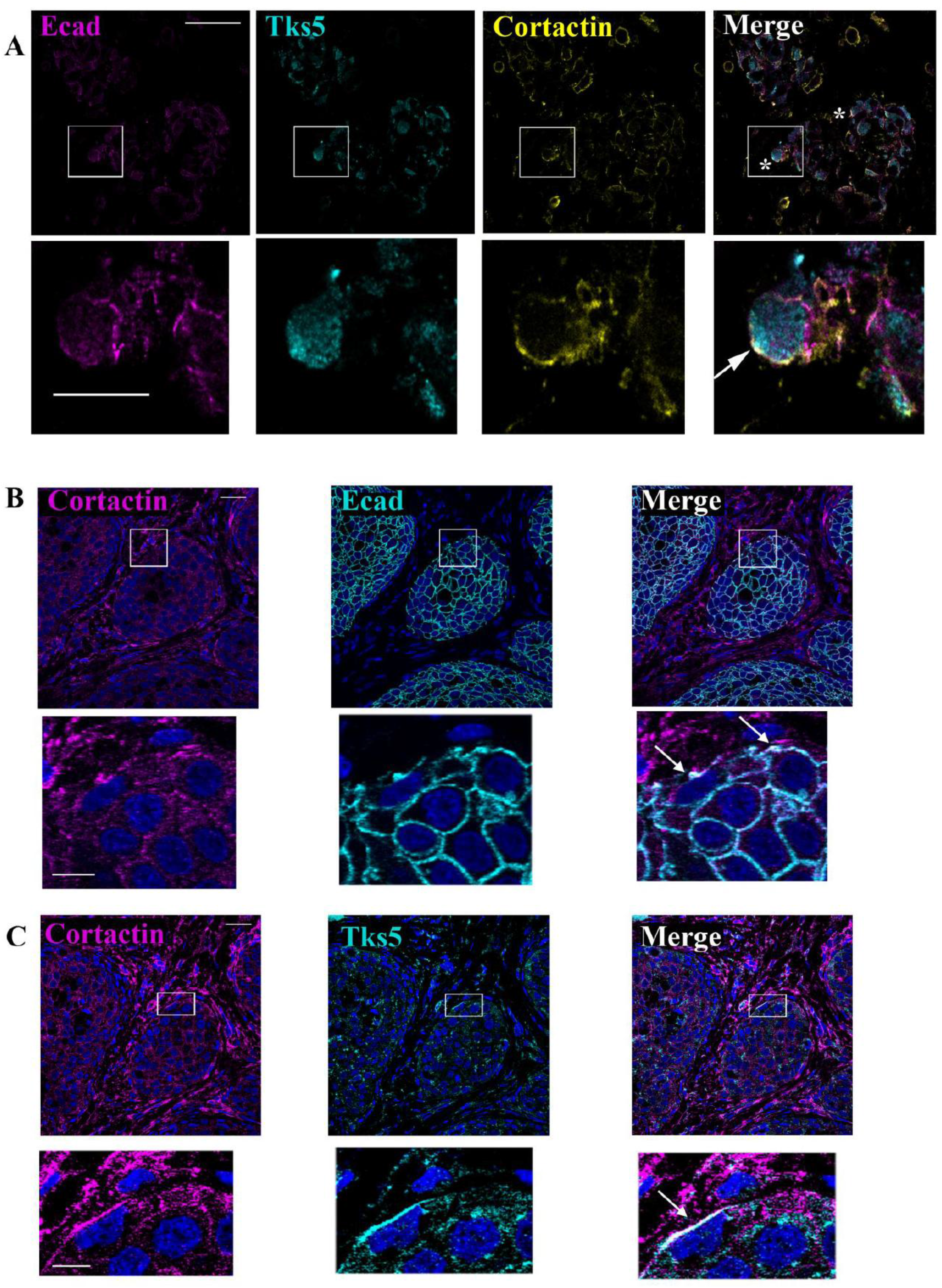
E-cadherin localizes in invadopodia-like structures *in vivo.* **(A)** Triple E-cadherin, Tks5, and Cortactin immunostaining in sections from patient tumours. White squares represent magnified views. White arrow indicates invadopodia containing E-Cadherin. Scale bars represent 10 µm (top panel) or 2 µm (magnification panel). The triple Cortactin/Tks5/E-cadherin colocalization was observed in 3 out of 6 patient tissues (n=6). **(B)** E-cadherin and Cortactin or **(C)** E-cadherin and Tks5 double immunostaining in serial sections from subcutaneous tumours of BxPC3 cells implanted in mice. Nuclei were stained using Dapi. White squares represent magnified views. White arrows indicate spots of Cortactin, Tks5 and E-cadherin colocalization. Scale bars represent 40 µm (top panels) or 10 µm (magnification panel).

Moreover, Cortactin or Tks5 were immunostained with E-cadherin in human pancreatic cancer BxPC-3 cells that had been ectopically implanted in mice. On serial tissues sections, colocalization of Cortactin/E-cadherin and Cortactin/Tks5 were observed at tumour cell plasma membranes in close contact with the microenvironment ***(Fig. 2B & 2C)***. Overlap coefficients are: <u>E-cadherin/cortactin (78.2%) and Tks5/cortactin (60.2%).</u>

Even if these structures are not common, these data confirm, in vivo, the existence of interactions between E-cadherin and all the invadopodia components. This suggests the localization of E-cadherin inside invadopodia-like-structures.

### E-cadherin interacts with MT1-MMP in invadopodia and is recycled through Rab7 and Rab11 pathways

To understand if this surprising localization of E-cadherin depends on a random distribution we first investigated the impact of cell-cell interaction on invadopodia formation. We observed that the number of invadopodia decreases in cells exhibiting intercellular contacts ***(Fig.3A)***, indicating a possible competition between the formation of cell-cell interactions and invadopodia. This suggests that the endocytic and exocytic fluxes of cell-cell contacts components are crucial for invadopodia activity.

**Fig.3:**
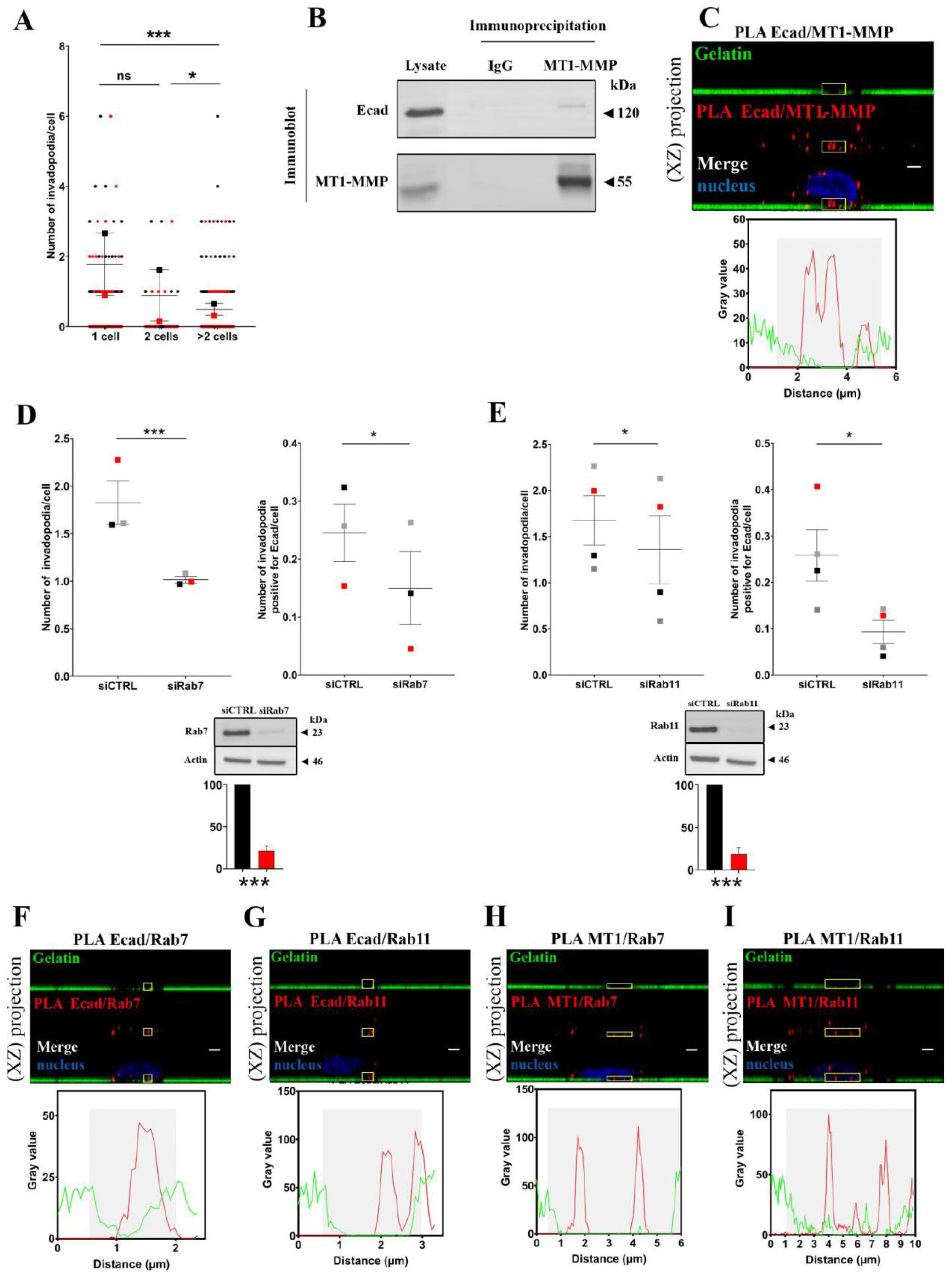
E-cadherin interacts with MT1-MMP in invadopodia and is recycled through Rab7 and Rab11 pathways. **(A)** Cell–cell interactions inhibited invadopodia formation. The number of invadopodia per cell was measured as described in Methods section. The graph represents the distribution of invadopodia in isolated cells (1 cell), cell doublet (2 cells) or groups superior of 2 cells (> 2 cells). Raw data are shown with coloured dots. Mean from 2 independent experiments are indicated with coloured squares. Errors bars represent mean ± SEM. n=2. **(B)** Equal amounts of BxPC-3 cell lysate were immunoprecipitated using either anti-MT1-MMP or non-specific (IgG) antibodies. After SDS-PAGE and transfer onto PVDF membrane, protein complexes were detected using anti-E-cadherin or anti-MT1-MMP antibodies. Control was performed using BxPC-3 lysates. A representative experiment of 3 (n=3) **(C)** E-cadherin and MT1-MMP colocalize inside invadopodia. After E-cadherin and MT1-MMP immunostaining, E-cadherin–MT1-MMP complexes were detected using PLA. Z-stack confocal acquisitions were performed. Top panel: The amplification spots (in red) localize in a gelatin-degradation area. Scale bar = 2 µm. Bottom panel: Fluorescence intensity quantification of the region of interest indicated by the yellow square on the top panel. The gelatin degradation area is identified in grey. A representative image of 2 experiments in triplicates with 3 acquisitions for each (n=2). **(D-E)** BxPC-3 cells were treated for 48h with siRNA control (siCTRL) or siRNA against **(D,)** Rab7 (siRab7) or **(E,)** Rab11 (siRab11) before invadopodia assay. **(D, E)** Left panel: Quantification of active invadopodia at the ventral surface of each cell. Right panel: Quantification of active invadopodia exhibiting E-cadherin per cell. Means from 3 **(D)** or 4 **(E)** independent experiments indicated with coloured squares. Errors bars represent mean ± SEM. Bottom panels: Equal amounts of cell lysate (25 µg) were subjected to SDS-PAGE, then transferred onto PVDF membrane. Graphs represent the mean ± SEM of Rab7 or Rab11 protein expression from 3 independent cell transfection. **(F)** E-cadherin and Rab7; **(G)** E-cadherin and Rab11; **(H)** MT1-MMP and Rab7; **(I)** MT1-MMP and Rab11. Z-stack confocal acquisitions were performed on fixed cells. Left panels: The amplification spots (red) localize with a degradation spot of the fluorescent gelatin (green). Scale bars represent 2 μm. Right panels: Fluorescence intensity quantification of the regions of interest indicated by the yellow square on the left panel. Images in 2D view for (**C**) and (**F-I**) are available in***Fig.S4B*** and negative control (PLA probe PLUS/MINUS) is available in ***Fig.S4D*.** **(F-I)):** A representative image of 2 experiments in triplicates with 3 acquisitions for each (n=2).

Vesicular transport has been shown to be crucial for invadopodia formation by facilitating the trafficking of MT1-MMP to the plasma membrane (Linder, 2015). On the other hand, E-cadherin undergoes cycles of endocytosis, sorting and recycling to the plasma membrane through Rab7 and Rab11 vesicles (Brüser & Bogdan, 2017; Terciolo *et al*, 2017). We therefore assessed if E-cadherin interacts with MT1-MMP and if its targeting to invadopodia depends on the same recycling process. Biochemical studies showed that a pool of E-cadherin co-precipitates with MT1-MMP, thus suggesting that these molecules can associate together ***(Fig.3B)***. Furthermore, PLA documented E-cadherin interaction with MT1-MMP in different cellular localizations: cell membrane, cytoplasmic vesicles and within invadopodia structures ***(Fig.3C)***. We found that both Rab7 ***(Fig.3D)*** or Rab11 ***(Fig.3E)*** depletion decrease the number of invadopodia containing E-cadherin. Moreover, E-cadherin-Rab7 ***(Fig.3F)*** and E-cadherin-Rab11 complexes ***(Fig.3G)*** were observed in several cytoplasmic vesicles some of which are localized in the immediate vicinity of the degradation areas. MT1-MMP was also detected in these compartments ***(Fig.3H & 3I)***. Altogether, these data indicate that E-cadherin is trafficked to invadopodia *via* an active recycling process through Rab7 and Rab11 pathways. They also demonstrate that MT1-MMP and E-cadherin could interact with each other inside invadopodia and are trafficked through the same pathway.

### E-cadherin adhesive activity is required for invadopodia formation

We next explored the role of E-cadherin in invadopodia formation and function using a cellular system engineered to silence E-cadherin expression by shRNA. Specifically, we generated stable BxPC-3 shEcad (E-cadherin depletion) and control BxPC-3 shCTRL (no cadherin depletion) cells (Siret *et al*, 2018). We found that E-cadherin silencing promotes a significant decrease in the number of cells forming invadopodia ***(Fig.4A)***. This was accompanied by a reduction of degradation areas ***(Fig.4B)*** and a significant decrease in the number of invasive structures per cell ***(Fig.4C)***. At the opposite, forced E-cadherin expression in E-cadherin deficient cells (***Fig.4D***) promotes (Dalle Vedove *et al*, 2019)a significant increase of degradation areas (***Fig.4D & 4E***).

**Fig.4:**
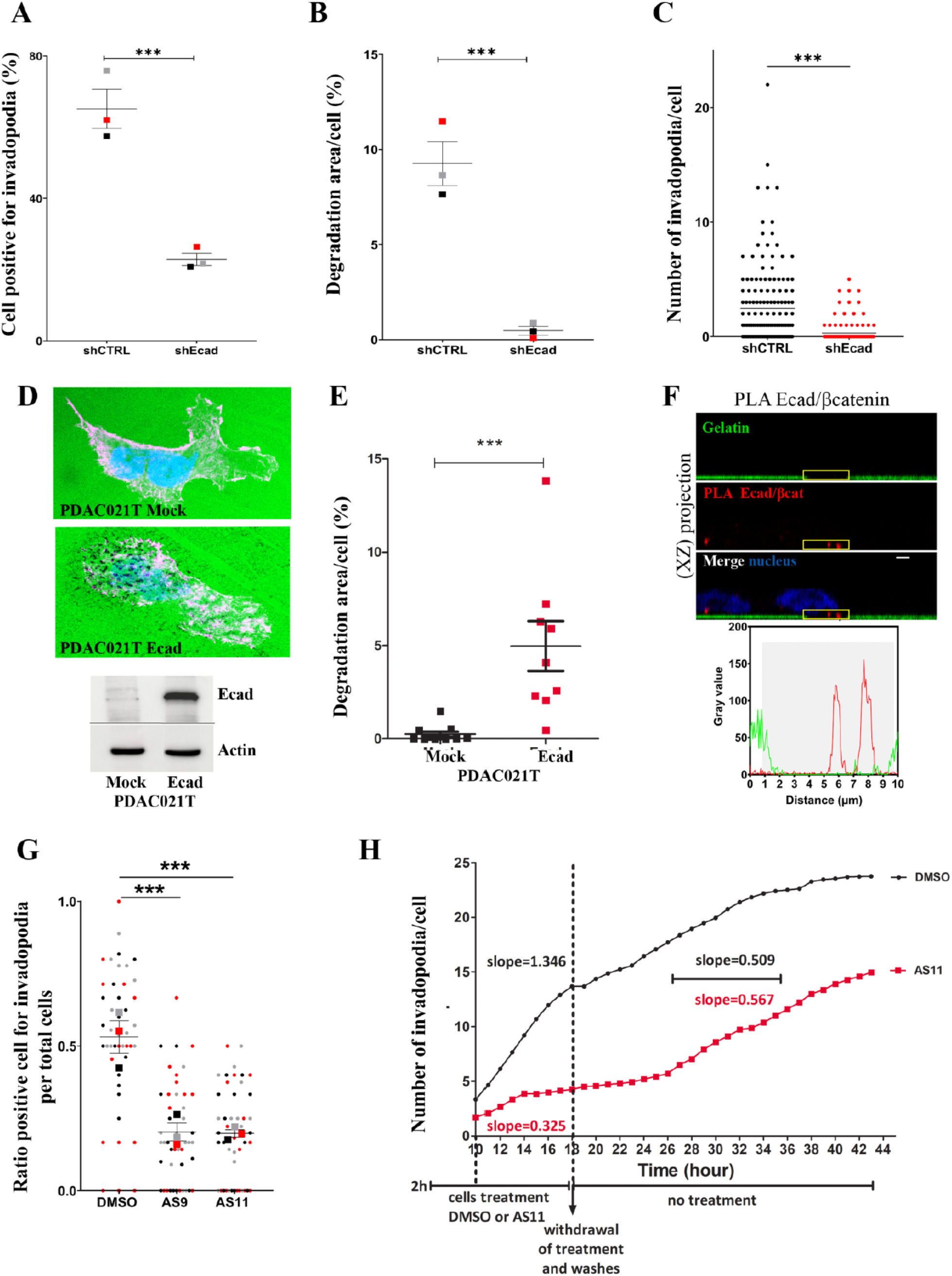
E-cadherin adhesive activity is required for invadopodia formation. **(A-C)** Invadopodia assays were performed using BxPC-3 control (shCTRL) and E-cadherin depleted cells (shEcad) cell lines. **(A)** The number of cells exhibiting active invadopodia were quantified. Means from 3 independent experiments are indicated with coloured squares. Errors bars represent Mean ± SEM. n=3 **(B)** The normalized gelatin degradation area at the ventral surface of the cells was evaluated. Means from 3 independent experiments are indicated with coloured squares. Errors bars represent mean ± SEM. n=3. **(C)** The distribution of the number of invadopodia per cell was determined A representative graph of 3 experiments (n=3). Images in 2D view for (**A**) are available in ***Fig.S4C*.** **(D-E)** Invadopodia assays were performed using PDAC021T Mock (no E-cadherin expression) and PDAC021T Ecad, (E-cadherin expression) cells. The E-cadherin expression was assessed by western blot.The normalized gelatin degradation area at the ventral surface of the cells were evaluated. Representative results from 3 independent experiments. **(F)** E-cadherin and β-catenin interact within invadopodia. E-cadherin–β-catenin complexes were detected using a PLA. Z-stack confocal acquisitions were performed. Top panel: the amplification spot (red) localizes in a degradation spot of FITC-labelled gelatin(green). Bottom panel: fluorescence intensity quantification of the region of interest indicated by the yellow square on the top panel. Scale bar represents 2 µm. A representative image of 2 experiments in triplicates with 3 acquisitions for each (n=2). Images in 2D view for (**F**) is available in ***Fig.S4D*.** **(G)** E-cadherin inhibition decreases invadopodia formation. Ratio of cells exhibiting active invadopodia in treated (AS9 or AS11) and untreated (DMSO) BxPC-3 cells were evaluated. Means from 3 independent experiments are indicated with coloured squares. Raw data are shown with coloured dots. Errors bars represent mean ± SEM. n=3. **(H)** Invadopodia assays were performed using BxPC-3 shCTRL. Cells were seeded for 2h on coverslips coated with FITC-labelled gelatin, then treated for 16h with DMSO or AS11. Cells were then washed and incubated in DMEM/10% fetal calf serum for an additional 24h period. Invadopodia formation was analysed by videomicroscopy by capturing images every hour, 8h after addition of the compounds. The number of gelatin degradation zones appearing just below the cell body is estimated for each hour. The graph is representative of an experiment carried out three times (n=3).

E-cadherin requires interactions with catenins to be functional (Mège & Ishiyama, 2017). PLA indicated that E-cadherin associates with β-catenin within invadopodia ***(Fig.4F)***. Moreover, we found that two synthetic E-cadherin inhibitors, AS9 and AS11, which block trans-interactions of E-cadherin molecules in junctional complexes (Dalle Vedove *et al*, 2019) reduced the number of cells exhibiting invadopodia ***(Fig.4G)***. By using videomicroscopy we analysed the impact of AS11 on the kinetic of invadopodia formation. AS11 decreased the rate of invadopodia appearance by 75% for the cells that still perform invadopodia, since the slope of the curves is 0.325 for AS11 treated cells versus 1.346 for control cells ***(Fig.4H*** *and **Fig.S2)***. The inhibitory effect of AS11 is rescued by removing the compounds. Indeed, cells resume invadopodia formation after 8h of latency with the same speed than control cells, when the E-cadherin inhibitor is removed. The slopes of the curves are 0,567 for the cells previously treated with AS11 and 0,509 for the control cells. These data confirmed the presence of a pool of functional E-cadherin at the invadopodial membrane and strongly suggest a role of E-cadherin in invadopodia structuring.

### An E-cadherin/Arp3 complex is detected into invadopodia

To determine the mechanisms by which E-cadherin expression could regulate the formation of invadopodia, we analysed by mass spectrometry the full proteome of E-cadherin depleted BxPC-3 cells (shEcad) versus cells expressing E-cadherin (shCTRL). We came up with a list of 64 proteins down-regulated and 80 proteins up-regulated when E-cadherin is depleted. The analysis of these data using the Ingenuity Pathway Analysis (IPA) software suggests at least 8 signalling pathways deregulated upon E-cadherin depletion, of which some could be crucial for invadopodia formation ***(Fig.5A and Fig. S3)***. Among enriched pathways, the actin nucleation by ARP/WASP complex was particularly interesting as this complex has been described in E-cadherin trafficking (Kovacs *et al*, 2002). As expected, interactome analysis identified the Arp2/3 complex as a partner of E-cadherin and β-catenin interaction networks ***(Fig.S3)***. We therefore focused on this pathway and found that E-cadherin depletion induced a down-regulation of all members of the Arp2/3 complex ***(Fig.5B)***. Western blot analysis additionally showed down-regulation of Arp3 subunit expression in E-cadherin-silenced cells ***(Fig.5C)***. Furthermore, PLA experiments indicate that Arp3 associates with both Cortactin and E-cadherin close to gelatin degradation areas, suggesting E-cadherin/Arp3 complex implication in invadopodia structuring ***(Fig.5D & 5E)***.

**Fig.5:**
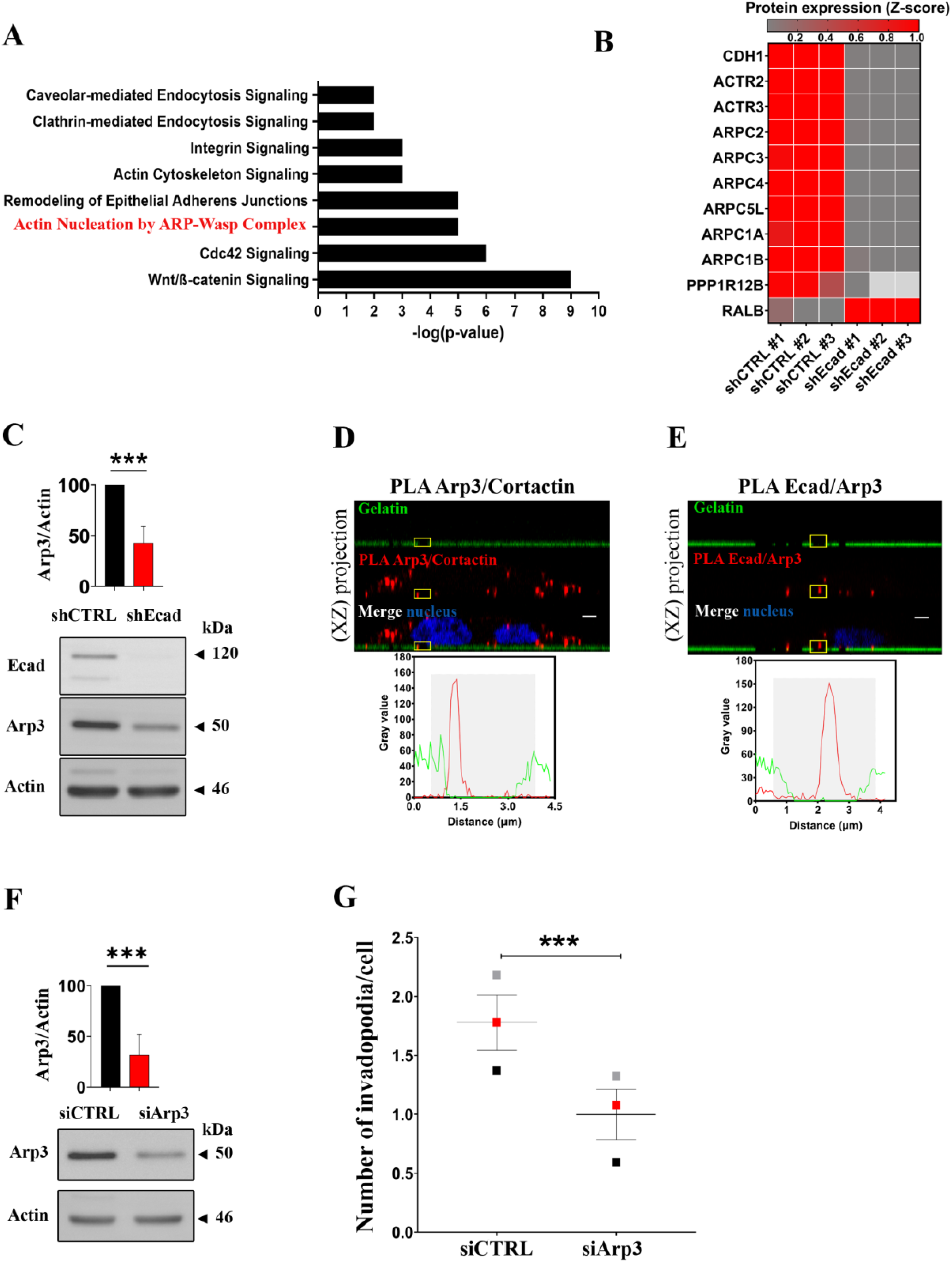
An E-cadherin/Arp3 complex is detected into invadopodia. **(A)** Most deregulated signalling pathways in BxPC-3 shEcad compared with BxPC-3 shCTRL cells as determined by IPA analysis of proteome data. The enrichment score on the graphic is represented by -log(p-value). n=3 **(B)** Heatmap of the Z-score of E-cadherin (CDH1) expression, RALB and proteins associated with actin nucleation through Arp2/3 complex pathway in BxPC3 shEcadh cells compared with BxPC3 shCTRL cells. Red and grey denote increase and decrease in protein expression, respectively. Three independent protein extractions were analysed (#1, #2, #3). n=3 **(C)** Western blot analysis of Arp3 expression in BxPC-3 shCTRL and shEcad cell lines. Equal amounts of cell lysate (25µg) were loaded on 8% polyacrylamide gel. After SDS-PAGE migration and transfer onto PVDF membrane, E-cadherin, Arp3 and actin were detected using specific antibodies. Bottom panel: representative western blot from 6 independent cells lysates. Top panel: Quantification of Arp3 expression from mean ± SEM. (n=6). **(D, E)** Protein–protein interactions in invadopodia revealed by PLA. **(D)** Arp3–Cortactin and **(E)** Arp3–E-cadherin interactions. Z-stack confocal acquisitions were performed on fixed cells. Top panels: The amplification spots (red) localize with a degradation spot of FITC-labelled gelatin (green). Cell nuclei are shown in blue. Bottom panels: Fluorescence intensity quantification of the region of interest indicated by the yellow square on the left panel. Scale bar represents 2µm. **(D):** A representative image of 2 experiments in triplicates with 3 acquisitions for each (n=2). **(E):** A representative image of 2 experiments in triplicates with 3 acquisitions for each (n=2). Images in 2D view for (**E**) are available in ***Fig.S4D*** and and negative control for (**D**-**E**) is available in ***Fig.S4D*.** **(F)** BxPC-3 cells were treated for 48h with control siRNA (siCTRL) or siRNA against the Arp3 subunit (siArp3). Arp3 protein expression in BxPC-3 cells treated by siCTRL or siArp3. Equal amounts of cell lysate (25µg) were subjected to SDS-PAGE, then transferred onto PVDF membrane. Arp3 and actin were detected using specific antibodies. The graph represents the mean ± SEM of Arp3 protein expression from 3 independent cell transfection. n=3. **(G)** After a treatment during 48h with control siRNA (siCTRL) or siRNA against the Arp3 subunit (siArp3) cells were plated for 16h onto FITC-labelled gelatin. The graph represents the quantification of active invadopodia formed per cell. Data corresponds to a mean from three independent experiments indicated with coloured squares. Errors bars represent mean ± SEM. n=3.

To functionally assess the implication of E-cadherin/Arp3 complex on invadopodia formation, we generated Arp3-silenced cells ***(Fig.5F)***. Reassuringly, we found that Arp3 depletion promoted a significant decrease in the number of invadopodia formed per cell ***(Fig.5G)***. Altogether, these data highlight the importance of the E-cadherin in the actin nucleation process through ARP/WASP complex. In the absence of the E-cadherin, Arp3 is down-regulated, and the actin nucleation does not take place, both preventing the formation of actin protrusions.

### E-cadherin is a structuring component of invadopodia

The association between E-cadherin and Arp3 promotes a signal for actin assembly during adherens junction formation (Kovacs *et al*, 2002). If the E-cadherin/Arp2/3 complex is involved in the structuring of invadopodia, this should be reflected in the observation of complex formation prior to matrix degradation.

In 12% of cells, both E-cadherin and Actin organizes into overlapping rings at the ventral cell surface prior gelatin degradation (***Fig.6A*** *step 1 and* ***Fig.6B***). If actin rings always associated with E-cadherin rings, the reverse is not true. This strongly suggest that E-cadherin ring structuration precedes actin assembly. Less frequently (3% of the cells) E-cadherin/Actin rings are associated with starting degradation area (***Fig.6A*** *step 2 and* ***Fig.6B***), indicating that these structures enriched in both E-cadherin and Actin represent invadopodia precursors. In 50% of the cells, large area of degradation accumulates with punctiform actin labelling associated or not with a E-cadherin staining. This suggests that after invadopodia maturation, E-cadherin dispersion precedes actin disassembly (***Fig.6A*** *step 3 and* ***Fig.6B***).

**Fig.6:**
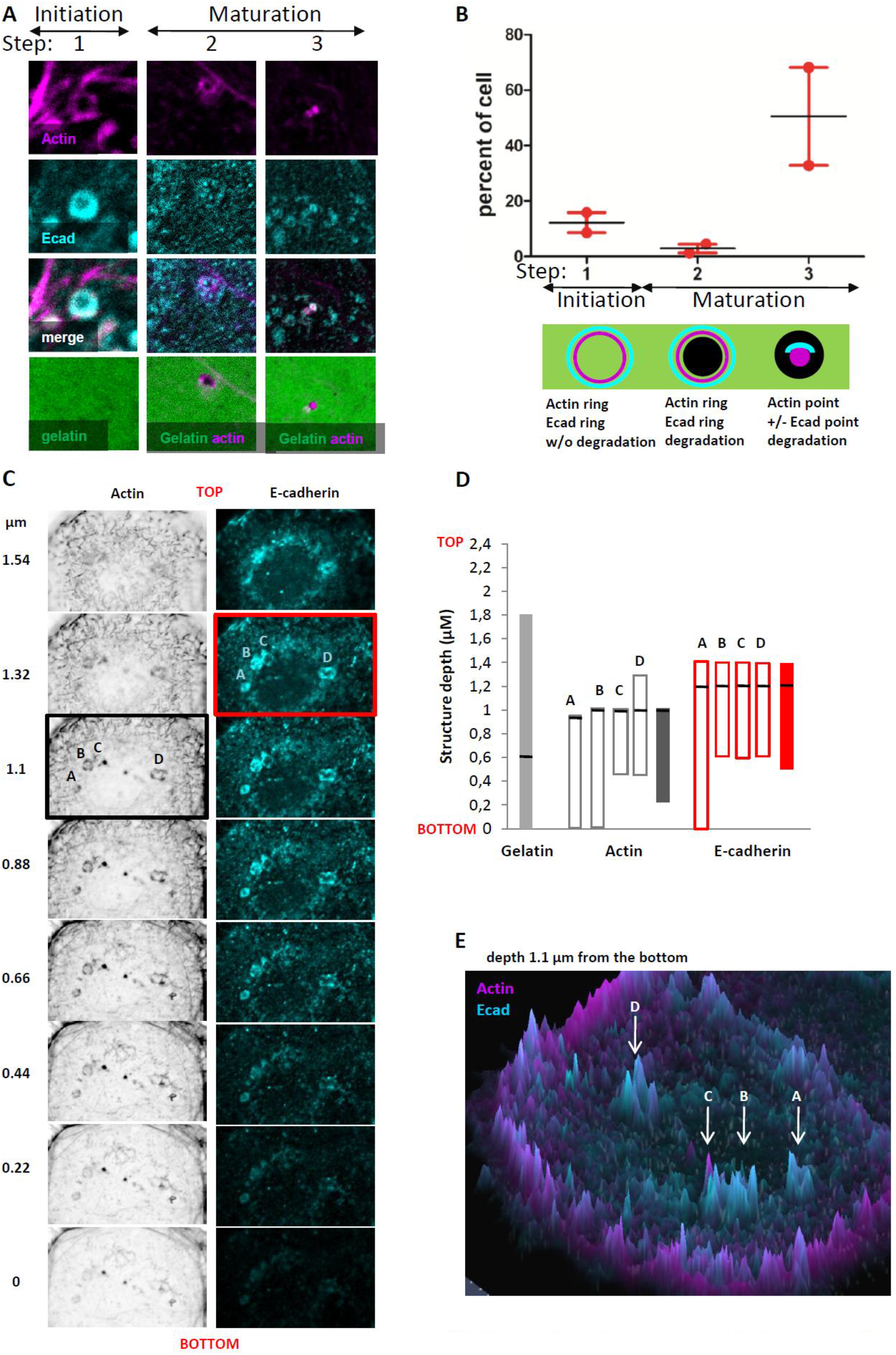
E-cadherin is a structuring component of invadopodia. Invadopodia assays were performed as previously described. (**A, B)** After confocal acquisition, analysis of structures evidence 3 kinds of invadopodia: step 1 initiation: In the absence of FITC-labelled gelatin degradation actin ring overlays with E-cadherin ring, step 2 maturation: spot of associated with both actin and E-cadherin ring, step3: FITC-labelled gelatin degradation associated with actin spot in the presence or absence of E-cadherin. (**B**), Percentage of cells showing these kinds of structure(n=2). (**C-F**) Images of Invadopodia assays were acquired using AiryScan module of Zeiss LSM 880 confocal microscope. Acquisitions were performed inside the gelatin sheet every 0.22µm. Bottom indicated the bottom detection of gelatin. In (**C)**, labelling of both Actin E-cadherin are given for each section. Intensity quantifications were performed using FIJI software and the maximal intensity for Actin and E-cadherin is mentioned by a black or red box, respectively. In (**D**) Z-labelling ranges are positioned with bars and the section with the more intense labelling is mentioned by a dark line. Dark bars represent the average of A, B, C, D structures. **(F)** Actin (magenta) and E-cadherin (cyan) staining detected 1,1 μm above the gelatin bottom using AiryScan module. Peaks represent labelling intensity of each molecule at this Z-section.

We observed that a part of E-cadherin ring is localized less deeply than actin ring in invadopodia (***Fig.6C, D and E***). Moreover, analysis of the localization of the maximal labelling intensity of the 2 molecules confirm this observation (***Fig.6C*** *and **D***).

Representation of the intensity of E-cadherin and Actin staining using Airyscan acquisitions, for a section taken 1.1µm from the bottom of the gelatin, shows that these rings are not blended into the background (***Fig 6E***).

Taken together, these results demonstrate a structuring role for E-cadherin during invadopodia formation.

## Discussion

E-cadherin was associated for years as a tumour suppressor. However, high levels of E-cadherin expression have been demonstrated in various invasive and metastatic cancer with epithelial traits suggesting that E-cadherin may have inefficient suppressive activity or even worst, could promote metastasis instead of suppressing tumour progression (Putzke *et al*, 2011; Padmanaban *et al*, 2019; Shen & Kang, 2019). Moreover, studies demonstrated that E-cadherin expression in E/M hybrid cells expressing E-cadherin might confer collective migratory ability to tumour cells, allowing them to survive during transit and colonization in distinct organs (Reichert *et al*, 2018; Shen & Kang, 2019).

To address the role of E-cadherin in PDAC aggressiveness, we modulated E-cadherin expression in pancreatic cell models and analysed the effect on cell invasion. According to our data, the contribution of E-cadherin on cancer cell invasion can be summarized as follows:

<u>E-cadherin is an early component of invadopodia.</u> (i) Originally localized in the adherens junctions, E-cadherin can be endocytosed and recycled back to the invadopodial membrane simultaneously with MT1-MMP. Both Rab7 and/or Rab11 vesicle-dependant pathways are required for this trafficking; (ii) Once translocated into the immature invadopodia, E-cadherin interacts with several components, such as Arp2/3 and Actin; <u>(iii) In association with Actin,</u> <u>E-cadherin forms a ring that precedes invadopodia degradative activity;</u> (iv) E-cadherin–β-catenin trans-interactions at invadopodial membrane suggest the establishment of new adherens-like junctions, allowing actin tension required for the protrusion scaffold ***(see graphical abstract*)**.

Invadopodia are hallmarks of various invasive cells (Yamaguchi, 2012; Meirson & Gil-Henn, 2018; Luo *et al*, 2021; Linder *et al*, 2023). They have been extensively studied in cell culture and have now been detected in *in situ* tissue explants, tissue sections and in vivo models (Génot & Gligorijevic, 2014; Lohmer *et al*, 2014; Chen *et al*, 2019). Invadopodia are supposed to represent promising therapeutic target to prevent cancer metastasis (Luo *et al*, 2021). Our ex vivo and in vivo results strengthen the physiological relevance of invadopodia in PDAC as the two most used invadopodia markers (Cortactin and Tks5) colocalize preferentially at cell plasma membranes in close contact with the extracellular matrix. According to this, we postulate that BxPC3 cell lines are a suitable research model for invadopodia studies in PDAC.

We provide multiple lines of evidence that E-cadherin is a key component of the invadopodial membrane. (1) E-cadherin localizes with both Cortactin and Tks5 close to the ECM surrounding tumour clusters in patient tissues; (2) a pool of E-cadherin but not P-cadherin is detected at the invadopodial membrane in the pancreatic BxPC-3 cell line, pancreatic cancer primary culture and a cell line derived from an inflammatory breast cancer. Moreover, E-cadherin interacts with the main components of invadopodia, including Cortactin, Tks5 and MT1-MMP. Furthermore, E-cadherin is distributed in invadopodia when cell–cell contacts are reduced. E-cadherin is trafficked to the invadopodial membrane, as MT1-MMP, through Rab7 and Rab11 recycling routes; (3) E-cadherin is found in purified fraction of invadopodia; (4) Modulation of E-cadherin expression is associated with invadopodia formation. Likewise, inactivation of E-cadherin trans-interaction in junctional complexes by drugs reversibly blocked invadopodia development, (5<u>) E-cadherin forms a ring that associates with actin ring</u> <u>to form an invadopodia precursor structure.</u> Some other compounds of the junctional complexes, including tight junctions (ZO-1), and Gap junctions (Connexin 43), may regulate invadopodia formation (Hu *et al*, 2018; Chepied *et al*, 2020). However, their ability to organize invadopodia has not been described.

The Arp2/3 complex polymerizes actin filaments as branches from existing filaments and powers various cell processes including cell motility, endocytosis, vesicle trafficking and adherens junction stability (Krause & Gautreau, 2014; Pandit *et al*, 2020). Its impact on actin polymerisation is critical for invadopodia-based invasion by driving cell protrusions through the ECM and maintaining tight apposition of surface-exposed MT1-MMP with the ECM (Monteiro *et al*, 2013). Here, three lines of evidence suggest a link between E-cadherin and Arp2/3 complex. First, IPA identified Arp2/3 as a direct partner of the E-cadherin interaction network. This is in agreement with studies that implicated Arp2/3 complex as a ke y actin assembly factor at E-cadherin-mediated cell–cell contacts (Kovacs *et al*, 2002). Second, E-cadherin associates with Arp3 in invadopodia. Finally, E-cadherin depletion induces a downregulation of the members of the Arp2/3 complex. Therefore, impaired actin nucleation, and E-cadherin depletion, prevents the formation of the actin protrusion which normally sustains invadopodia.

From these results we postulate that E-cadherin allows the establishment of membrane junctions in invadopodia structures. Cadherins have been described as participating in the formation of junctions within a single cell. For instance CDHR5 and CDHR2, indirectly anchored with the core actin bundle, are known to orchestrate microvillus crosslinking in intestinal enterocytes (Dooley *et al*, 2022).

E-cadherin localization in invadopodia requires intracellular trafficking, including endocytosis and recycling via Rab7 or Rab11 vesicles dependent pathways. <u>However,</u> <u>pathways involved in invadopodia activity, including Rab2A dependent vesicles and exocyst</u> <u>complex (Sakurai-Yageta *et al*, 2008; Kajiho *et al*, 2018) may also be involved in E-cadherin</u> <u>trafficking to invadopodia. Further works are needed to define exactly how E-cadherin is</u> <u>transported to invadopodia.</u>

To summarize, we demonstrated that E-cadherin promotes pancreatic cancer cell invasion by regulating invadopodia formation. The proinvasive function of E-cadherin and its related signalling mechanism need to be further explored. Importantly, these findings open new avenues towards uncovering innovative options for earlier diagnosis and anti-invasive therapy of pancreatic cancer.

## Additional informations

AD, VR and FA conceived and designated the study and the experiments. AD, SG, VR, RB, FA, FS performed the experiments, AD, VR and SG analysed the data. SA performed the proteomic analysis, PS, IJ and ND provide inputs of the study. AD, VR and FA wrote the manuscript.

## Acknowledgements

The authors thank Philippe Chavrier (Institut Curie, Paris, France) for providing helpful advice at the beginning of the project, Magalie Benard (PRIMACEN Rouen, France) Sylvie Thuault and Eric Mas (CRCM) for discussions. We thank Flavio Maina and Avais Daulat for their feedbacks and suggestions on the written manuscript. We thank Magda Rodrigues (CRCM Misc platform) for support and advice. Proteomic analyses were performed at the mass spectrometry facility of Marseille Proteomics supported by IBISA (Infrastructure Biologie Santé et Agronomie), Plateforme Technologique Aix-Marseille, Cancéropole PACA, Région Sud-Provence-Alpes-Côte d‘Azur, Fonds Européen de Développement Régional (FEDER) and Plan Cancer. We are grateful to the ICEP (IPC/CRCM experimental pathology) core-facility for histological processing of tumour samples.

## Funding information

This work was supported by INCa (Grants number 2018-078 and 2018-079), Cancéropôle PACA, DGOS (labellisation SIRIC), Amidex Foundation, Ligue contre le Cancer, Fondation de France and INSERM.

## Data availability

The mass spectrometry proteomics data have been deposited to the ProteomeXchange Consortium (http://www.proteomexchange.org) via the PRIDE partner repository with the dataset identifier PRIDE: PXD017895.

## Competing Interests

The authors declare no conflict of interest.

## Additional Data 1

### Mass spectrometry analysis and data processing protocol

Proteomes from E-cadherin depleted cells (BxPC-3 shEcad) were compared to control cells (BxPC-3 shCTRL) by label-free quantitative mass spectrometry analysis. 15 µg of each cell lysate was loaded on NuPAGE 4-12% Bis-Tris acrylamide gels (Life Technologies) to stack proteins in a single band that was stained with Imperial Blue (Thermo Fisher Scientific) and cut from the gel. Gel pieces were submitted to an in-gel trypsin digestion (Shevchenko *et al*, 1996). Peptides were extracted from the gel and dried under vacuum. Samples were reconstituted with 0.1% trifluoroacetic acid in 4% acetonitrile and analyzed by liquid chromatography (LC)-tandem mass spectrometry (MS/MS) using an Orbitrap Fusion Lumos Tribrid Mass Spectrometer (Thermo Electron, Bremen, Germany) with a nanoRSLC Ultimate 3000 chromatography system (Dionex, Sunnyvale, CA). Peptides were separated on a Thermo Scientific Acclaim PepMap RSLC C18 column (2 µm, 100A, 75 µm × 50 cm). For peptide ionization in the EASY-Spray nanosource in front of the Orbitrap Fusion Lumos Tribrid Mass Spectrometer, spray voltage was set at 2.2 kV and the capillary temperature at 275 °C. The Orbitrap Lumos was used in data-dependent mode to switch consistently between MS and MS/MS. The time between master scans was set to 3 seconds. MS spectra were acquired with the Orbitrap in the range of m/z 400–1600 at a FWHM resolution of 120,000 measured at 400 m/z. AGC target was set at 4.0e5 with a 50 ms maximum injection time. For internal mass calibration, the 445.120025 ions were used as lock mass. The more abundant precursor ions were selected, and collision-induced dissociation fragmentation was performed in the ion trap to have maximum sensitivity and yield a maximum amount of MS/MS data. Number of precursor ions was automatically defined along run in 3 s windows using the ―Inject Ions for All Available parallelizable time option‖ with a maximum injection time of 300 ms. The signal threshold for an MS/MS event was set to 5,000 counts. Charge state screening was enabled to exclude precursors with 0 and 1 charge states. Dynamic exclusion was enabled with a repeat count of 1 and duration of 60 s.

Relative intensity-based label-free quantification (LFQ) was processed using the MaxLFQ algorithm from the freely available MaxQuant computational proteomics platform, version 1.6.3.4. Spectra were searched against the human database extracted from UniProt on the 1^st^ September 2020, which produced 20,375 entries (reviewed). The false discovery rate (FDR) at the peptide and protein levels were set to 1% and determined by searching a reverse database. For protein grouping, all proteins that could not be distinguished on the basis of their identified peptides were assembled into a single entry according to the MaxQuant rules. Statistical analysis was done with Perseus program (version 1.6.14.0) from the MaxQuant environment (www.maxquant.org). Quantifiable proteins were defined as those detected in above 70% of samples in one condition or more. To obtain a normal distribution, protein LFQ normalized intensities transformed using base 2 logs. Missing values were replaced using data imputation by randomly selecting from a normal distribution centred on the lower edge of the intensity values that simulates signals of low abundant proteins using default parameters (a downshift of 1.8 standard deviation (s.d.) and a width of 0.3 of the original distribution). To determine whether a given detected protein was specifically differential, a two-sample *t*-test was done using permutation-based FDR-controlled at 0.01 and employing 250 permutations. The *p* value was adjusted using a scaling factor s0 with a value of 0.4. Analysis was done on biological triplicates, each run three times on mass spectrometers. The mass spectrometry proteomics data have been deposited to the ProteomeXchange Consortium via the PRIDE (Perez-Riverol *et al*, 2019) partner repository with the dataset identifier PXD021795.

## Legends to supplementary figures

**Fig.S1:**
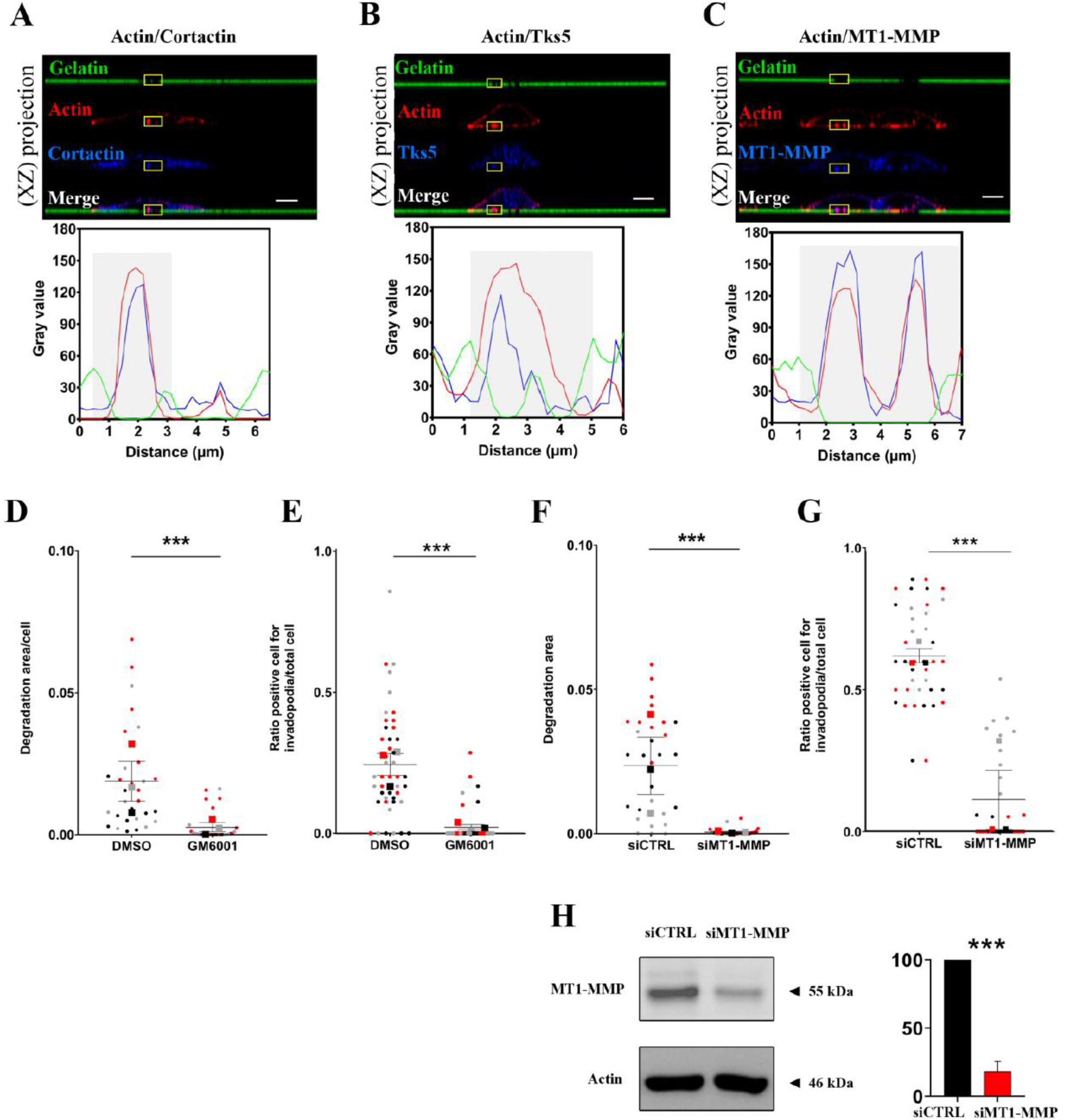
Invadopodia characterization in BxPC-3. **(A-C)** BxPC-3 cells were plated for 16h onto FITC-labelled gelatin then fixed. Actin, Tks5 and MT1-MMP were immunostained. **(A)** Actin (red) and Cortactin (blue), **(B)** Actin (red) and Tks5 (blue), **(C)** Actin (red) and MT1-MMP (blue). Z-stack acquisitions were then performed. Top panels: Colocalization of actin spots with a degradation zone of the gelatin (black spot) represents active invadopodia. Scale bar = 2 µm. Bottom panels: Fluorescence intensity quantification of the region of interest indicated by the yellow square on the left panel. The gelatin degradation area is identified in grey. **(A-C)** A representative image of 5 experiments with 3 acquisitions for each (n=5). Images in 2D view for (**A-C**) are available in ***Fig.S4A*** **(D)** Quantification of gelatin degradation area at the ventral surface of treated (GM6001 inhibitor) and control (DMSO-treated) BxPC-3 cells. **(E)** Ratio of positive cells for active invadopodia in treated (GM6001 inhibitor) and control (DMSO-treated) BxPC-3 cells. **(F)** Quantification of gelatin degradation area at the ventral surface of siCTRL or siMT1-MMP treated cells. **(G)** Ratio of cells exhibiting active invadopodia in siCTRL and siMT1-MMP treated cells… **(D-G)** 10 microscopic fields are quantified for each condition of the 3 experiments; Mean from 3 independent experiments are indicated with coloured squares Errors bars represent Mean ± SEM **(H)** Western blot analysis of MT1-MMP protein expression in BxPC-3 siCTRL and siMT1-MMP cells. BxPC-3 cells were treated for 48h with siRNA control (siCTRL) or siRNA against MT1-MMP (siMT1-MMP). MT1-MMP and actin were detected using specific antibodies. The graph represents the mean ± SEM from three independent cell transfections. n=3

**Fig.S2:**
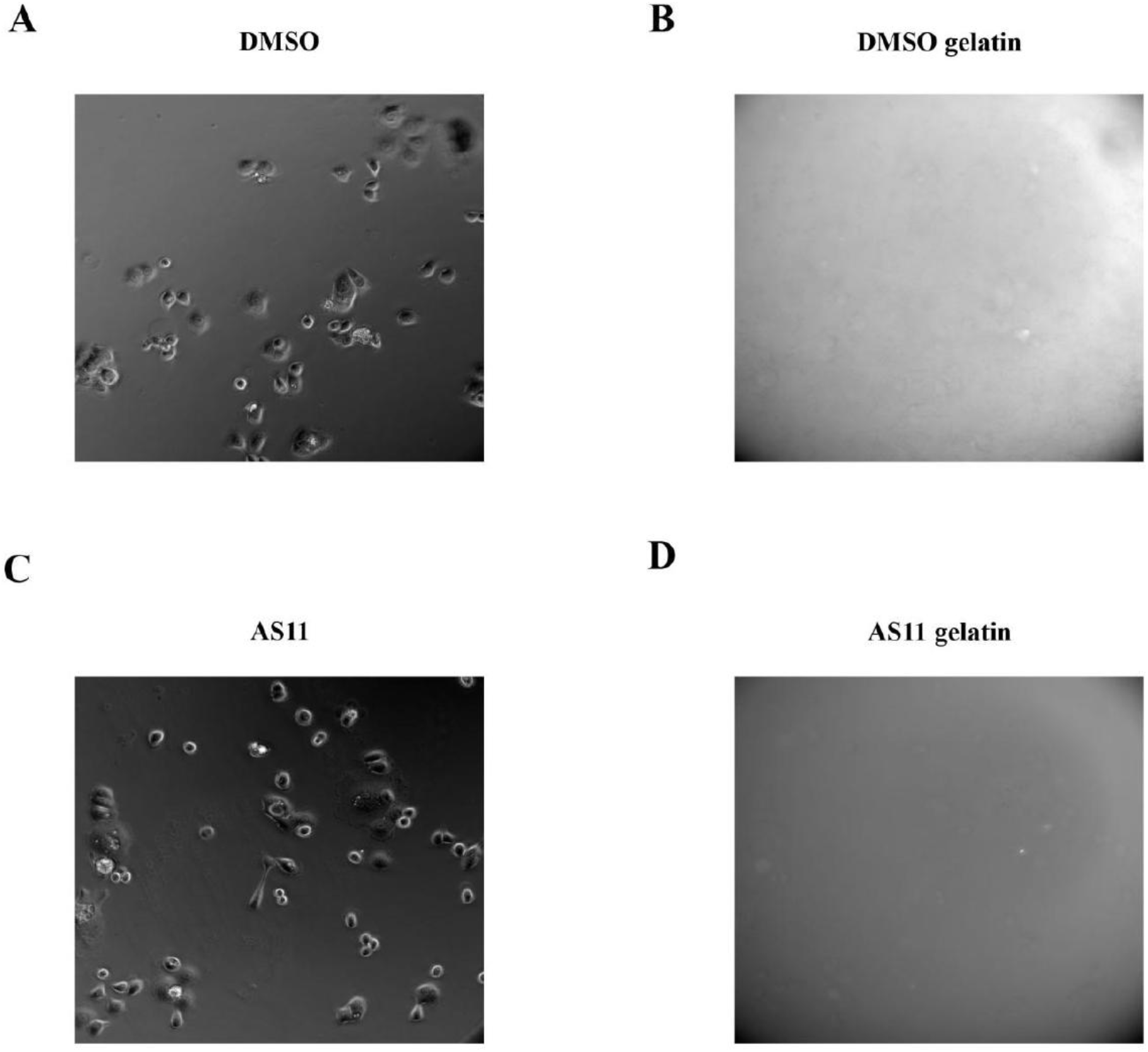
videos for invadopodia dynamic. Invadopodia assays were performed using BxPC-3 shCTRL. Cells were seeded for 2h on coverslips coated with FITC-conjugated gelatin, then treated for 16h with DMSO (**A** and **B**) or AS11 (**C** and **D**). Cells were then washed and incubated in DMEM/10% fetal calf serum for an additional 24h period. Invadopodia formation was analysed by videomicroscopy by capturing images every hour, 8h after addition of the compounds. **A** and **C** represent bright field images; **B** and **D** represent gelatin degradation areas. The number of gelatin degradation zones appearing just below the cell body is estimated for each hour. The graph (representative of an experiment carried out three times) is available in Fig. 4H

**Fig.S3:**
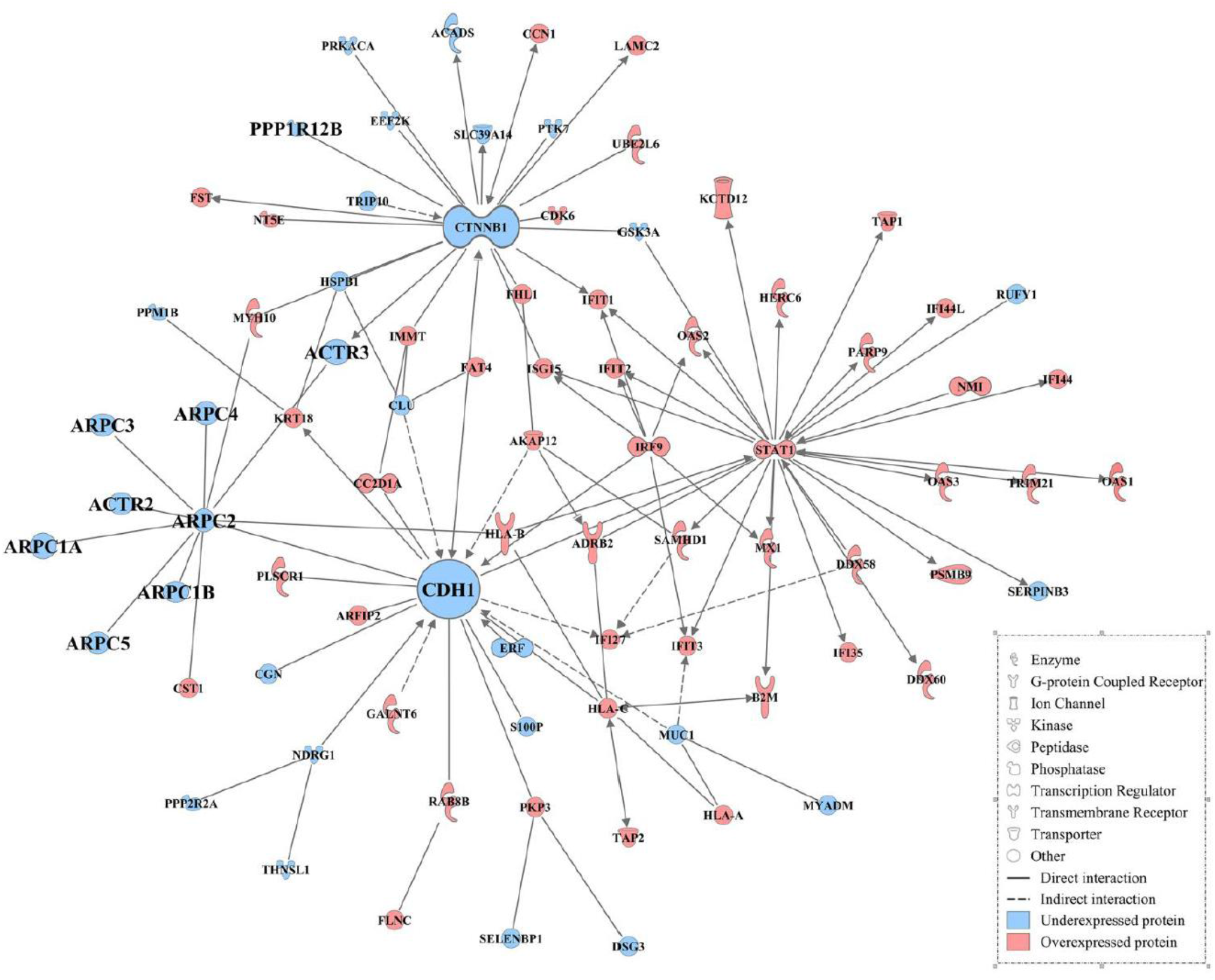
Central interactome for E-cadherin. Proteins identified by mass spectrometry were analysed using Ingenuity Pathway Analysis (IPA). The central interactome for E-cadherin was built with IPA software showing deregulated proteins identified by mass spectrometry.

**Fig.S4:**
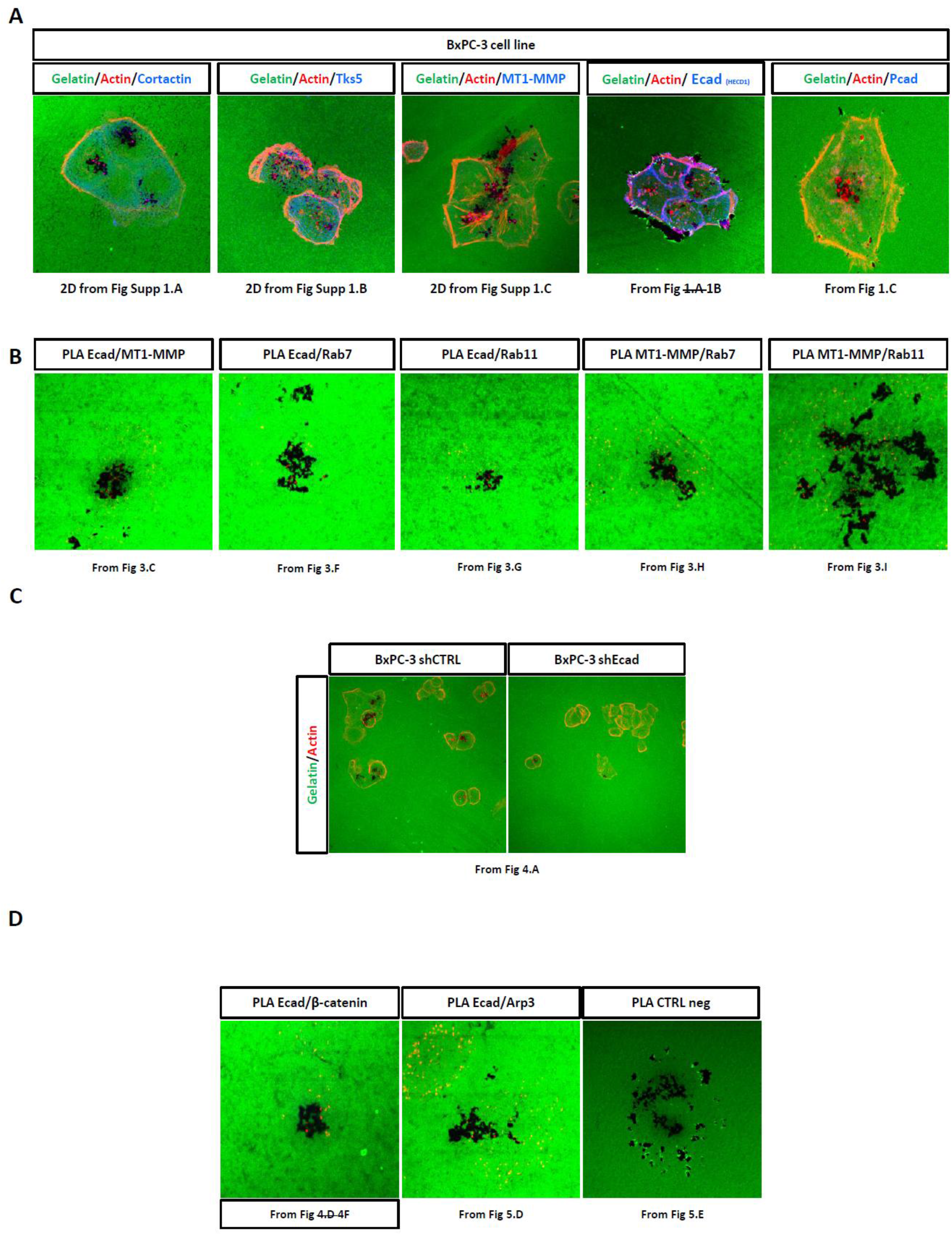
Controls for invadopodia imagery. (**A**): 2D view of invadopodia labelling presented in Fig. 1 and Fig. S1. (**B**): 2D view of invadopodia labelling presented in Fig. 3. (**C**): 2D view of invadopodia labelling presented in Fig. 4A. (**D**): 2D view of PLA labelling (presented in Fig. 4) and negative control of PLA (presented in Fig. 3 and 5).

## Notes

### Competing Interest Statement

The authors have declared no competing interest.

### Summary of Updates

Revisions process allow us to demonstrate a strucurant action of E-cadherin for invadopodia formation.

